# Temperature fluctuations influence predictions of landscape-scale spruce budworm defoliation in a physiologically-informed species distribution model

**DOI:** 10.1101/2024.08.19.608715

**Authors:** Emily N. Black, Deepa S. Pureswaran, Katie E. Marshall

## Abstract

**Aim:** While many studies on ectotherm thermal tolerance consider temperature exposure, the frequency of temperature exposures is emerging as an important and generally overlooked driver of survival and fitness which may influence species ranges. We use a physiologically-informed species distribution model to evaluate the influence of temperature fluctuations on the historical distribution and intensity of defoliation of a Lepidopteran forest pest, and on predicted future defoliation.

**Location:** Eastern Canada.

**Time period:** 2006-2016, projections to 2041-2070.

**Major taxa studied:** *Choristoneura fumiferana* (Lepidoptera: Tortricidae, spruce budworm).

**Methods:** We combined publicly-available maps of spruce budworm-induced defoliation between 2006-2016 in the Canadian province of Quebec with climate, forest composition, and de novo temperature fluctuation predictors to train a species distribution model. Our model evaluated how predictor categories influence spruce budworm defoliation and compared these results to a model trained without temperature fluctuations. Additionally, we predicted future spruce budworm defoliation under 2041-2070 climate change conditions using the models trained with and without temperature fluctuation predictors to determine the impact of temperature fluctuations on future defoliation predictions.

**Results:** We found that the inclusion of temperature fluctuation predictors improved model performance, and these predictors ranked highly in importance relative to predictors in other categories. The model trained with temperature fluctuation predictors also predicted vastly different defoliation distribution and severity across Quebec, Ontario, and Labrador than the model trained without them under future climate.

**Main conclusions:** Our study reveals the previously overlooked importance of temperature fluctuations on landscape-scale spruce budworm defoliation and demonstrates the importance of including physiologically-informed predictors in species distribution models. It also provides a novel framework for including thermal variation in correlative species distribution models of ectotherms.

## Introduction

Temperature directly affects ectotherm survival, growth, metabolism, and fecundity, and is a major factor determining species’ spatial distributions (Régnière, 2009; Régnière, Powell, et al., 2012). In temperate regions, the type and intensity of thermal stress varies dramatically across the year. In winter, ectotherms may not be able to forage (Findsen et al., 2014), and risk tissue damage from freezing (Storey & Storey, 1989). Temperate ectotherms can also be highly sensitive to fall and spring conditions (Sinclair, 2015) as unfavourable temperatures in these seasons can deplete energy stores, reduce survival, and induce non-lethal costs such as limited fecundity (Marshall & Sinclair, 2015; Roe et al., 2024). High temperatures in summer may induce elevated metabolic rates and macromolecular damage, which incur survival and fitness costs (González-Tokman et al., 2020). Given the significant effects of temperature on ectotherm fitness, is important to study how physiological adaptations to temperature contribute to environmental responses and to species distributions.

While many physiological and ecological studies use mean and extreme temperatures to characterize thermal regimes across the range of ectotherms, these do not reflect conditions in the wild where temperature fluctuates on multiple time scales (Dillon et al., 2016). While organisms can experience temperature fluctuations on many frequencies, the predominant periods are yearly (seasonal) and daily (Dillon et al., 2016), and these fluctuations can have significant impacts on fitness (Kern et al., 2015). Patterns of temperature fluctuations are expected to shift due to climate change, as Wang & Dillon (2014) found that global temperature variability has increased under climate change, and that this effect is particularly pronounced in temperate and polar regions. However, this varies by temperature threshold and season, as the frequency of extreme cold temperatures (below −20°C) and freezing temperatures (below 0°C) are expected to decrease in the boreal forest in 2081-2100 climate conditions (Contosta et al., 2024).

Fluctuations at different temperatures, seasons, and life stages can have varying and overlapping impacts on organismal physiology and fitness. This has been observed in a variety of taxa including fish, aquatic invertebrates, and lizards (Dickerson & and Vinyard, 1999; Dong et al., 2006; Kern et al., 2015; Putnam & Edmunds, 2011; Raynal et al., 2022); however, it is particularly well-characterized in insects (Marshall & Sinclair, 2012). Frequent temperature fluctuations within an insect’s optimal thermal range can positively impact insect fitness and fecundity by decreasing energy requirements for growth (Carrington et al., 2013). Previous exposure to high temperatures in the pea leafminer (*Liriomyza huidobrensis*) leads to higher expression of heat shock proteins and improved thermotolerance in subsequent heat exposures (Huang et al., 2007). At below-freezing temperatures, temperature fluctuations can positively or negatively affect insect fitness depending on the species and effect being studied (Marshall & Sinclair, 2012). For example, the frequency of cold temperature exposures is more detrimental than cold temperature duration or intensity to overwintering survival of (eastern) spruce budworm *Choristoneura fumiferana* (Lepidoptera: Tortricidae; Marshall and Sinclair 2015). Fluctuations reducing cold temperatures may also allow for repair of cold-damaged tissues during winter (Colinet et al., 2006), or by contrast, lead to de-acclimation of insects to winter conditions that increases the chance of chilling injury (Sobek-Swant et al., 2012), and therefore positively or negatively affect survival (Marshall & Sinclair, 2010, 2012, 2015). Overall, temperature fluctuations have a wide range of impacts on insect physiology, survival, and fitness (Marshall & Sinclair, 2012), and their effects may shift as climate change progresses.

Despite advances in our understanding of how temperature fluctuations drive physiology and fitness, these are not incorporated in methods for investigating species distributions despite their potential ability to improve predictions of species ranges (Maino et al., 2016). Species distribution models generally rely on associations between species distributions and macroenvironmental predictors including temperature means and extremes (aka “correlative” models). While these temperature metrics can suggest insights into species range drivers, they lack physiological grounding and interpretability (Kearney & Porter, 2009) and may not accurately predict species responses to climate change (Maino et al., 2016). Incorporating physiologically-informed predictors into correlative species distribution models enables them to directly incorporate drivers of fitness in addition to a suite of environmental predictors when predicting species ranges, and these models can make physiologically-informed predictions of species ranges without the significant data requirements of alternative methods such as mechanistic and biophysical models (Enriquez-Urzelai et al., 2019). Organisms for which there are extensive distribution data and well-characterized physiological mechanisms which drive fitness make excellent model species to implement these models and evaluate how the incorporation of physiologically-informed predictors into correlative species distribution models impacts the ability to predict and evaluate species ranges.

The spruce budworm is a major North American insect pest of boreal forests from Alaska to Newfoundland (Gray, 2008; Marshall & Roe, 2021; Natural Resources Canada, 2021). Females lay eggs in late summer, then first instar larvae hatch and develop into second instars that spin hibernacula and overwinter in trees until early May. After overwintering, larvae feed voraciously on coniferous trees before developing into adults. Outbreaks of this pest occur every 30-40 years and cause landscape scale defoliation: the larvae will eat expanding new buds of host trees (balsam fir, white spruce, black spruce; Mattson et al., 1991) in significant quantities. Several years of consecutive defoliation eventually weakens and kills the trees (Gray, 2008). A recent outbreak in Quebec resulted in 14.3 million hectares of forest damage via defoliation since 2006 (Ministère des Ressources naturelles et des Forêts, 2024). Outbreaks are potentially preventable if “hotspots” where populations are increasing are identified in the early stages (Johns et al., 2019; MacLean et al., 2019), which requires significant effort to understand drivers of outbreaks and defoliation severity, extensive forest monitoring, and prediction of future defoliation hotspots (Johns et al., 2019). Extensive aerial monitoring by provincial governmental agencies records the location and severity of spruce budworm defoliation across Canada, and this provincial scale lends itself well to the generation of spruce budworm species and defoliation distribution models (Ministère des Ressources naturelles et des Forêts, 2023; Natural Resources Canada, 2021).

Predictive distribution models of spruce budworm outbreaks have evaluated the importance of different environmental factors, including temperature, on outbreak likelihood and defoliation risk on a landscape scale (Bouchard & Auger, 2013; Gray, 2008; Li et al., 2020) and have predicted the effects of climate change on outbreak distribution and severity (Candau & Fleming, 2005; Johns et al., 2019; Régnière, St-Amant, et al., 2012). In particular, Gray (2008) found that patterns in outbreak distribution were partially explained by climate variables and forest composition, and that outbreaks were expected to be longer and more severe under climate change. Candau & Fleming (2005) also found that the distribution of spruce budworm outbreaks is strongly correlated with temperature: in particular, winter maximum and minimum temperature. However, these models generally use monthly, seasonal, and annual temperature means and extremes, which generally do not reflect conditions experienced by spruce budworm and may not be physiologically relevant. Temperature fluctuations affects the survival of spruce budworm in laboratory experiments (Marshall & Sinclair, 2015). Thus, using spruce budworm as a model species we determine if incorporating temperature fluctuations into species distribution models improves performance, and determine their importance relative to environmental factors such as host tree distribution and other climatic conditions.

Here, we provide a novel framework for incorporating thermal variation into a species distribution model using the spruce budworm *(Choristoneura fumiferana*) as a model system. Based on previous work that demonstrated the importance of temperature fluctuations for spruce budworm survival (Marshall and Sinclair 2015), we hypothesized that temperature fluctuations influence the landscape-scale distribution of spruce budworm defoliation. Using historical records of defoliation collected in Quebec, we trained a spatial random forest regression model to predict the severity of historical defoliation using three selected general predictor categories (climate, forest, temperature fluctuations), and then determined the effect of including temperature fluctuations in the model on predicted defoliation under a 2041-2070 climate change scenario. We found clear evidence that temperature fluctuations influence the landscape-scale distribution of spruce budworm defoliation, and inclusion of these predictors significantly impact predictions of spruce budworm defoliation under climate change. This framework demonstrates the usefulness of incorporating physiological predictors into species distribution models, and may be useful for models of other ectotherm taxa whose survival and fitness is influenced by temperature fluctuations.

## Methods

### i) Collecting and aggregating spruce budworm defoliation data

The Ministère des Ressources naturelles et des Forêts in Quebec conducts annual aerial flyovers of forests to record damage due to major insect pests, the effects of disturbances such as forest fires, and the effectiveness of current pest control measures (Ministère des Ressources naturelles et des Forêts, 2024). Survey locations are based on the locations of insect damage from the previous year, surveys of diapausing larvae, and predictive modelling of future outbreaks (Ministère des Ressources naturelles et des Forêts, 2023, 2024). The location and severity of spruce budworm *(Choristoneura fumiferana*) defoliation patches are recorded during this survey (Ministère des Ressources naturelles et des Forêts, 2024). Spruce budworm defoliation patches collected from 1992-2006 (n_polygon_=4,958), 2007-2013 (n_polygon_=54,867), and 2014-2021 (n_polygon_=262,826) are stored as polygons in a publicly available file (see supp. table 1 for links to data sources). The polygons have the following attributes: year of defoliation, defoliation severity level and score, hectares of defoliation, Quebec administrative region, and the geometric shape of the defoliation polygon. Defoliation severity scores were 1 – mild canopy damage, 2 – moderate canopy damage, and 3 – severe canopy damage. Maps of Ontario defoliation were also collected, but were not used in subsequent analyses due to difficulties standardizing with the Quebec data (supp. methods, Figure S1).

We aggregated defoliation polygons for defoliation on a raster grid using the *terra* package (Hijmans et al., 2021; Figure 1) to model cumulative spruce budworm defoliation from the start of the most recent outbreak. While year-over-year species distribution modelling approaches have been used to evaluate spruce budworm outbreaks (see Srivastava et al., 2024), we chose to evaluate overall outbreak propensity of the environment by summarizing defoliation intensity across all years studied. These summarized values also reduce the influence of biotic factors on outbreak dynamics explored by our model, which vary significantly across space and time over the course of an outbreak (Bouchard & Auger, 2013; Royama, 1984). We mapped the presence and absence of defoliation, and the cumulative defoliation severity of each grid cell (i.e., how susceptible it was to spruce budworm damage) by summing the annual defoliation severity score of all polygons covering the centroid of a grid cell and assigning that value to the cell. The raster of spruce budworm defoliation in Quebec covered an area from −79.57°W to −61.55°W, and 45.42°N to 51.73°N. Grids had an initial resolution of approximately 412m × 412m; however, grids used in analysis were aggregated to a resolution of 300 arcsec (approximately 6.5km × 6.5km) to match predictor resolution.

**Figure 1.**
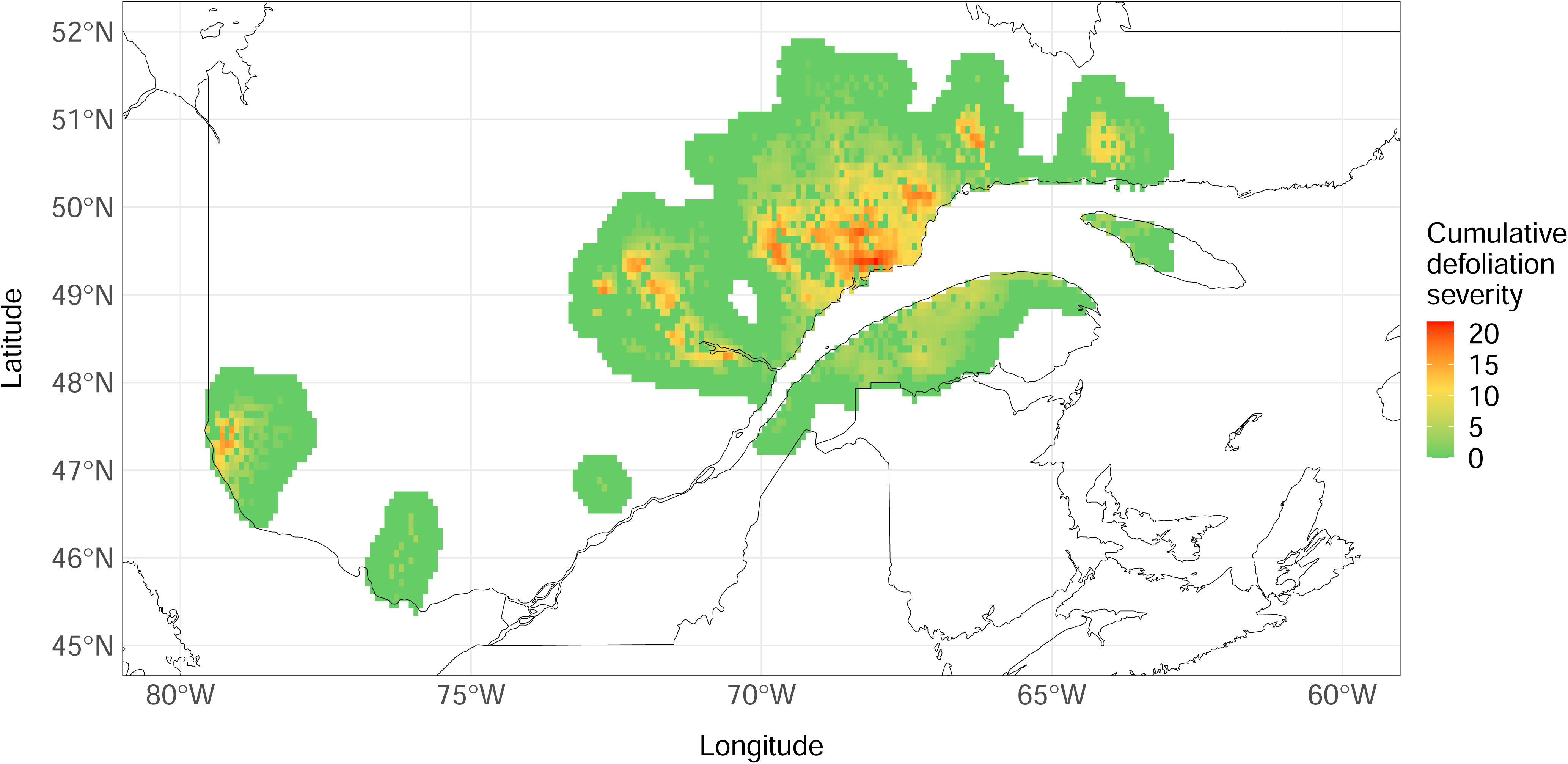
Distribution of spruce budworm cumulative defoliation severity across Quebec from 2006-2016, collected from aerial surveys conducted by *Ministère des Ressources Naturelles et des Forêts* in Quebec. Cumulative defoliation severity calculated by summing annual defoliation severity levels in each year from 2006-2016. Cells where defoliation is present have a mean cumulative defoliation severity of 6.63 ± 4.27. An additional 35km buffer with no measured defoliation was added to the mapped defoliation to serve as absences. n = 4,146 grid cells (total area = 233,269km^2^). Resolution: 6.5km × 6.5km.

Government records of spruce budworm defoliation indicate that the most recent outbreak in Eastern Canada began in 2006 (Natural Resources Canada, 2021); thus, we used defoliation patches from this year onwards in our analysis. To confirm that defoliation polygons aligned with known outbreak years, we plotted the cumulative amount of defoliation in each grid cell in which defoliation was observed between 1997 and 2021. Defoliation increased exponentially across Quebec 2006 onwards (Figure S2). While the outbreak beginning in 2006 is ongoing (Natural Resources Canada, 2021), high-resolution daily climate records were only available until 2016. Thus, we only included defoliation from 2006-2016 in analyses.

### ii) Collecting and aggregating predictor data

We chose to evaluate the effect of four general predictor categories on defoliation presence and severity: forest, climate, degree days, and temperature fluctuations. For a list of data sources, see supp. table 1; for a list of all predictors and their descriptions and assigned predictor categories, see supp. table 2.

For forest predictors, we obtained raster grids of the forest composition of Canada, including the distribution of spruce budworm host trees (white spruce, black spruce, balsam fir) and percent tree cover from the National Forest Inventory (NFI), a collaborative effort by federal, provincial, and territorial government agencies to generate continuous records of forest composition across Canada (*National Forest Inventory*, 2021). All grids had an initial resolution of 250m × 250m, and were upscaled to approximately 6.5km × 6.5km using bilinear interpolation (Hijmans et al., 2021).

For climate, temperature fluctuation, and degree day predictors, we obtained minimum and maximum daily temperatures and daily precipitation from 2006-2016 across eastern Canada from the CHELSA (Climatologies at High Resolution for the Earth’s Land Surface Areas) dataset. CHELSA calculated global daily fine-resolution temperature and precipitation values up to 2016 using downscaling of atmospheric temperatures and climatologies (Karger et al., 2017, 2021). Although many high-resolution climate products exist for our region of interest (e.g. T. Wang et al., 2016), we selected CHELSA primarily due to its generation of global daily temperatures which allowed us to generate accurate daily temperature fluctuation counts across a broad geographic region, as well as its generation of global monthly climate variables under several climate change scenarios. Additionally, CHELSA’s global distribution allows our methods to be more easily extrapolated to other species and regions of interest across the globe. From the daily CHELSA temperature values, we calculated the annual mean temperature, annual mean minimum and maximum temperature, and the absolute annual minimum and maximum temperature. We calculated these values for each month of the year, averaged across 2006-2016. We also obtained daily precipitation values from the CHELSA dataset to calculate the mean yearly and monthly precipitation. For the degree day predictors, using the CHELSA data we summed the difference between mean daily temperatures and 0°C annually and in each month, averaged across 2006-2016. Maps of historical climate and degree day data covered an area from −83.57°W to −57.55°W, and 41.42°N to 55.73°N, and had an initial resolution of 300 arcsec (approximately 6.5km × 6.5km).

To obtain temperature fluctuation predictors across the landscape, we calculated repeated temperature exposures, or the number of times the temperature in each grid cell crossed a particular temperature threshold daily each month (for example, the number of times the temperature crossed 0°C daily in January) based on temperature treatments performed by Marshall and Sinclair on 2nd instar spruce budworm (2015) and predominant periods of temperature fluctuations identified by Dillon et al. (2016). To calculate this, we interpolated the daily minimum and maximum temperatures from CHELSA to simulate change in daily temperature across an entire year at each grid cell. Then, we summed the number of times the temperature at each point crossed above a particular threshold (e.g., 0°C, −10°C) to calculate the frequency of cold exposure. We then averaged these values across 2006-2016 and transformed the points into a raster map with a resolution of 300 arcsec (approximately 6.5km × 6.5km). Overall, we generated a total of 169 climate, forest, and temperature fluctuation predictors (see supp. table 2 for description of all predictors).

To predict defoliation presence and severity in the future, we re-generated model predictors under simulated climate change conditions from 2041-2070. Since tree species in the eastern Canadian boreal forest show limited migration ability under climate change conditions (Boisvert-Marsh et al., 2022) and the majority of reductions in forest area under SSPs climate scenarios are expected to occur outside of North America (Popp et al., 2017), we used the same tree distributions as the historical models (*National Forest Inventory*, 2021).

To generate climate, degree day, and temperature fluctuation variables under climate change conditions, we used the monthly CHELSA CMIP5-predicted climatologies for 2041-2070 (Karger et al., 2017). These represent monthly temperatures in an example year under climate change scenarios, and were generated using the Geographic Fluid Dynamics Laboratory Earth System 4 (GFDL-ESM4; 10). We selected climatologies from the SSP370 climate scenario, which predicts a moderate to high amount of future global emissions (*The SSP Scenarios*, n.d.). While the SSP370 climate scenario includes substantial land-use and aerosol emission changes in addition to increased greenhouse gas emissions, these have a limited effect on projected solar radiation and precipitation in Eastern Canada as compared to other SSP scenarios (Shiogama et al., 2023). Additionally, the majority of land-use changes are projected to occur outside of North America (Popp et al., 2017). To simulate daily temperatures under this climate change scenario, we used the *chillR* package (Luedeling et al., 2023) to generate daily stochasticity in the monthly CHELSA climate change projections. The *chillR* package uses the CHELSA monthly projections to guide future climatic trends in the stochastic data, and historical daily temperatures from 2010-2016 CHELSA data to generate stochastic daily minimum and maximum temperatures that follow annual temperature trends (Figure S3). From the *chillR* daily stochastic weather output, we generated annual and monthly climatic predictors, degree day predictors, and temperature fluctuation predictors under SSP370 climate change conditions. As we were predicting into the future and the range of the eastern spruce budworm is expected to shift (Gray, 2008; Régnière, St-Amant, et al., 2012), we increased the spatial extent of the predictors to include −96.76°W to −59.51°W and 43.42°N to 53.75649°N. All predictors had a spatial resolution of approximately 6.5km × 6.5km.

### iii) Building and evaluating distribution models of defoliation

Though spruce budworm is found endemically through much of Canada’s boreal forest (Marshall & Roe, 2021), only certain populations will transition to “epidemic” (or outbreaking) dynamics with densities high enough to cause measurable defoliation. Thus, “absences” in our model are defined as areas where no outbreaks were observed and populations remain at low endemic densities rather than areas where spruce budworm are completely absent. We marked cells up to 35km away from the nearest point with defoliation as regions where defoliation was absent, creating a “buffer region” around the outbreak. Tree cover did not differ significantly between areas where defoliation was present and the buffer region, so no additional filtering of absence cells was performed. We also ensured grid cells used for training the models were within Quebec provincial boundaries. We used points within this buffer as true absences of defoliation in the model.

Exploratory modelling showed that random forest regression had the highest r-squared value (best preliminary performance) amongst several machine learning algorithms tested; thus, it was selected for further modelling analysis. To generate honest and spatially unbiased metrics of model performance, we used spatial cross-validation to reduce the effect of spatial autocorrelation on model performance and evaluation. To do this, we used the *spatialRF* package (Wright & Ziegler, 2017), which can generate spatially-explicit random forest models, evaluate the contributions of predictors locally and to overall model performance, and evaluate model performance using spatial cross-validation. We chose to evaluate the influence of temperature fluctuation predictors on defoliation using a random forest regression model predicting cumulative defoliation severity (i.e. summed severity scores).

Climate, degree day, forest, and temperature fluctuation predictors were highly correlated with one another, and therefore all predictors initially showed a high degree of overfitting and resulted in poor overall performance. Thus, we ran an initial model with all 169 predictors and default *spatialRF* hyperparameters (Wright & Ziegler, 2017) and obtained the ranked relative importance of each predictor. Then, we kept predictors with a high relative importance in the initial model, and eliminated less-important predictors with a correlation coefficient over 0.95 and subsequently a variance inflation factor (VIF) higher than 7.5 as compared to higher-ranking predictors (i.e. if two predictors are highly correlated with one another, the predictor with a higher importance to the initial model is kept while the other is removed; Zhang et al., 2023). These values are more conservative than those used in similar models (Dormann et al., 2013); however, due to the large number of variables which were highly correlated with one another, we decided more restrictive predictor selection was needed. This significantly reduced the number of predictors in the final model with all predictor categories from 169 to 12 predictors and the model trained without fluctuations from 96 to 8 predictors. Degree day predictors were not selected by this process; thus, they were absent from subsequent analysis. As some host tree predictors were not selected during predictor pruning, they were added back in to ensure predictions of defoliation under climate change were limited by the distribution of suitable food and habitat (which are not expected to change significantly under climate change scenarios; Boisvert-Marsh et al., 2022). We then re-tuned model hyperparameters using the *spatialRF* function “rf_tuning”.

To evaluate the model and determine the importance of selected predictors, we performed spatial cross-validation with a 4:1 training:testing data split to measure the root-mean-square error (RMSE) of the regression model, and the permuted increase in model error when individual predictors were removed (Wright & Ziegler, 2017). For evaluation and climate change projections, the model predicted the cumulative defoliation severity score. To determine the effect of temperature fluctuation predictors on model performance we built and evaluated a second model which did not include temperature fluctuation predictors. Predictor pruning using variance inflation factor was performed on climate and forest predictors alone using the same protocol as the initial model, and model hyperparameters were re-tuned for this model to ensure differences in performance were not due to poorly-adapted model hyperparameters (Wright & Ziegler, 2017).

To determine where the model performed more poorly, we investigated whether model error (residuals) varied predictably across the region. We also generated partial dependence plots to visualize the relationships between predictors and defoliation severity. Partial dependence plots show the relationship between the value of a predictor and the value of the predicted defoliation propensity generated by the model. This can be used to infer the effect of the predictor on spruce budworm defoliation.

### iv) Predicting defoliation under 2040-2071 climate change conditions

We used the random forest regression to predict regions across Quebec and Ontario (from −96.76°W to −59.51°W and 43.42°N to 53.75649°N) which may be at risk of defoliation in a simulated 2040-2071 climate scenario. We provided the model with the CHELSA CMIP5 climatology predictors and existing host tree distributions and had the model predict defoliation across the region. Then, to determine the effect of temperature fluctuations on predictions of defoliation under climate change, we re-predicted defoliation under climate change conditions using the model trained without temperature fluctuation predictors. To measure the effect of temperature fluctuations, we report deviations in cell-by-cell predictions between the full model and the model with temperature fluctuation predictors removed and investigated differences in the distribution of predicted defoliation.

All analyses were performed using R version 4.3.0 (R Core Team, 2021).

## Results

### i) Overall model performance

Overall, the regression model predicted cumulative defoliation severity with a root-mean-squared error (RMSE) of 2.64 ± 0.32 (SD) during evaluation (indicating predictions tended to deviate from actual defoliation severity by a value of 2.64). The model trained without temperature fluctuation variables had a RMSE of 3.01 ± 0.45 (SD), indicating model performance decreased when trained without temperature fluctuation predictors.

To understand where the regression model performed more poorly, we investigated the residuals of the regression model along a gradient of defoliation severity. We tested both a polynomial and linear fit and found that model error increased along a gradient of defoliation severity when considering grid cells where defoliation was present (linear regression fit: R^2^ = 0.37, F_1,1479_ = 866.3, *p <*0.001; Figure 2). While the relationship between residuals and defoliation severity was fit similarly by a polynomial and linear regression (ΔAIC = 0.55), we selected the simpler linear model. Grid cells where no defoliation was observed showed a wide range of residuals, but generally defoliation absence was predicted accurately (y-axis histogram, Figure 2B).

**Figure 2.**
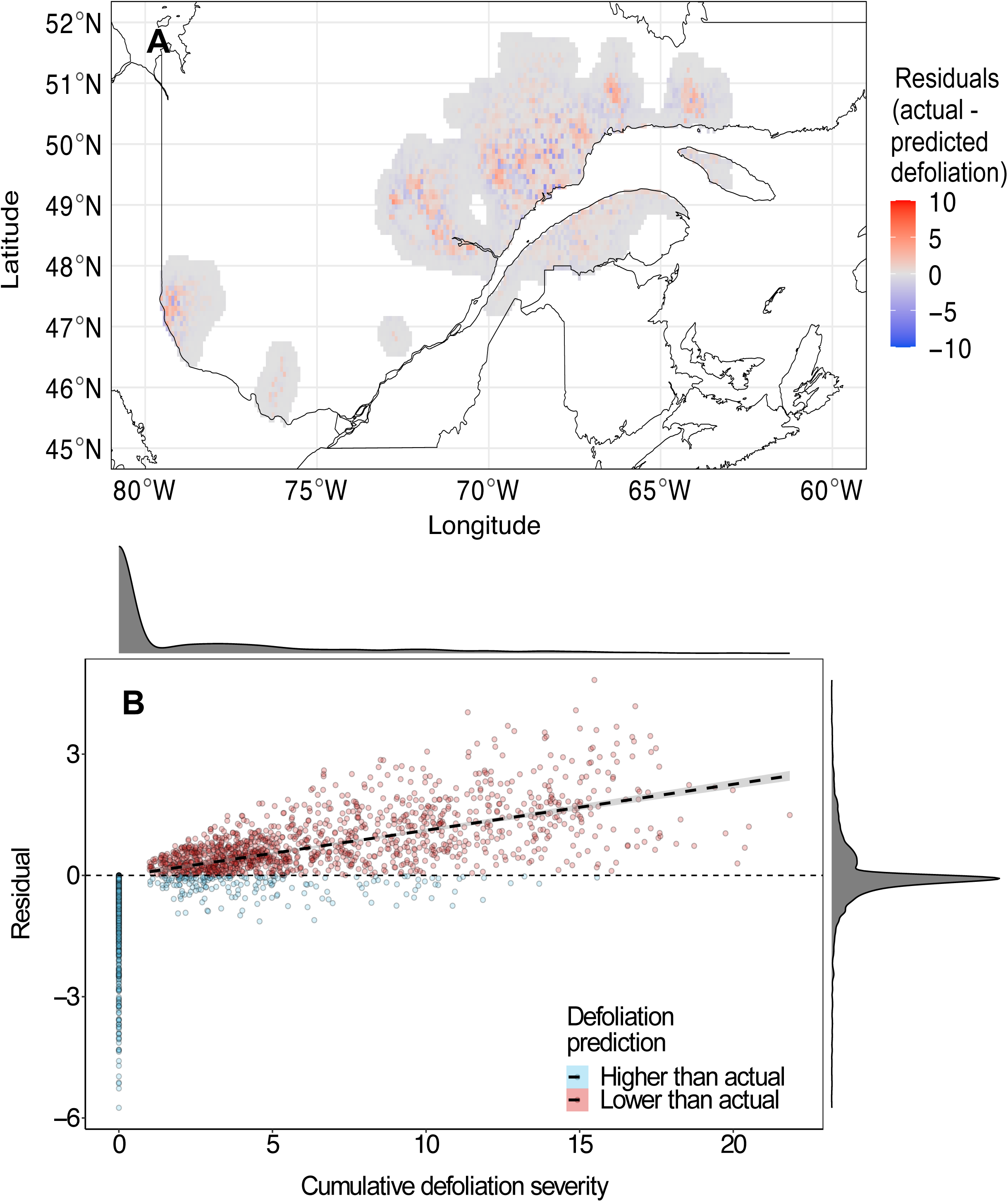
Regression model error tends to increase in areas with higher observed defoliation severity. **A)** Distribution of residuals of random forest regression model. Red cells indicate model underestimated defoliation severity, blue cells indicate model overestimated defoliation severity. **B)** Relationship between model residuals and actual values of cumulative defoliation severity. Histograms in margins show density of data across x- and y-axis values. Linear regression line only fit to cumulative defoliation severity (x-axis) values higher than zero (where defoliation was present). n_present_ = 1,481 grid cells.

### ii) Predictor contributions

Climate and temperature fluctuation predictors had the highest overall importance in the random forest regression model, with a mean increase in error (RMSE) when permuted of 1.95 ± 0.21 (SD) and 1.83 ± 0.35 (SD), respectively (Figure 3). Forest composition predictors had lower overall importance, with a mean increase in error (RMSE) when permuted of 1.48 ± 0.20 (SD). The distribution of local (cell-by-cell) importance of the top four most important variables also vary among predictors, with some showing higher importance localized to areas where defoliation is present or absent. For example, August mean precipitation generally had higher importance in areas where defoliation was present, while January −5°C fluctuations was generally more important in areas where defoliation was absent (Figure S4). Temperature fluctuation predictors make up six of 15 predictors in the model and rank relatively highly among other predictors. Five out of six temperature fluctuation predictors rank higher than all tree distribution predictors. Ranked predictor importance for models trained with and without temperature fluctuations are available in supp. tables 3A and 3B, respectively.

**Figure 3.**
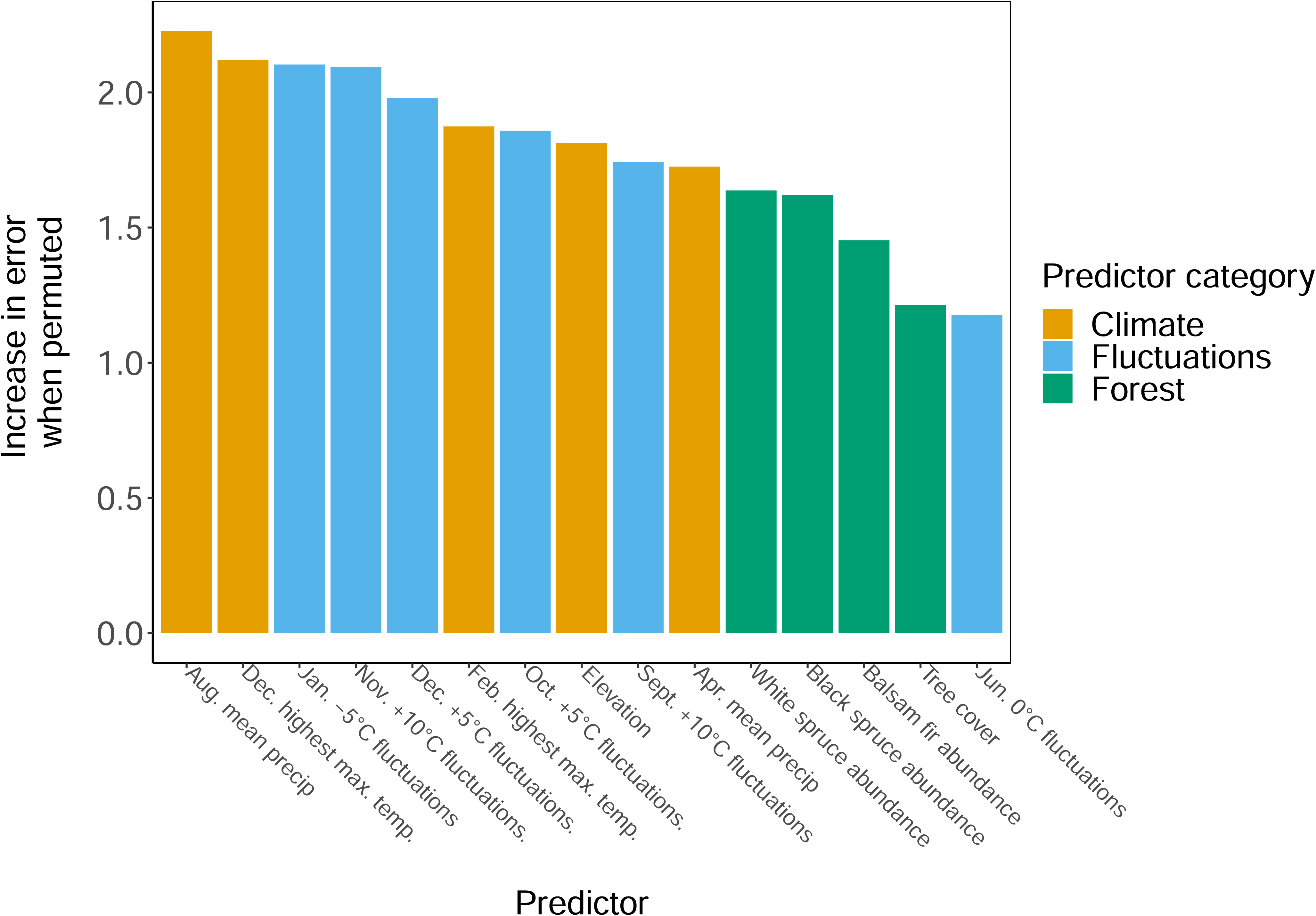
Climate and temperature fluctuation predictors are generally more important than forest variables in the random forest regression model. Bars represent increase in overall model error when predictor was removed from training model.

The relationships between cumulative defoliation severity and model predictors in the regression model are complex and depend on the quantiles of other predictors (Figure 4). Generally, defoliation is more severe with lower August mean precipitation and higher April mean precipitation. Defoliation severity also increases at lower elevations and higher tree cover (although relationships with specific tree species are variable). There is generally no consistent relationship between defoliation severity and fluctuation predictors, as these vary across months, temperatures, and quantiles of other predictors (Figure S5 shows relationship between values of the four most important predictors and observed and predicted defoliation).

**Figure 4.**
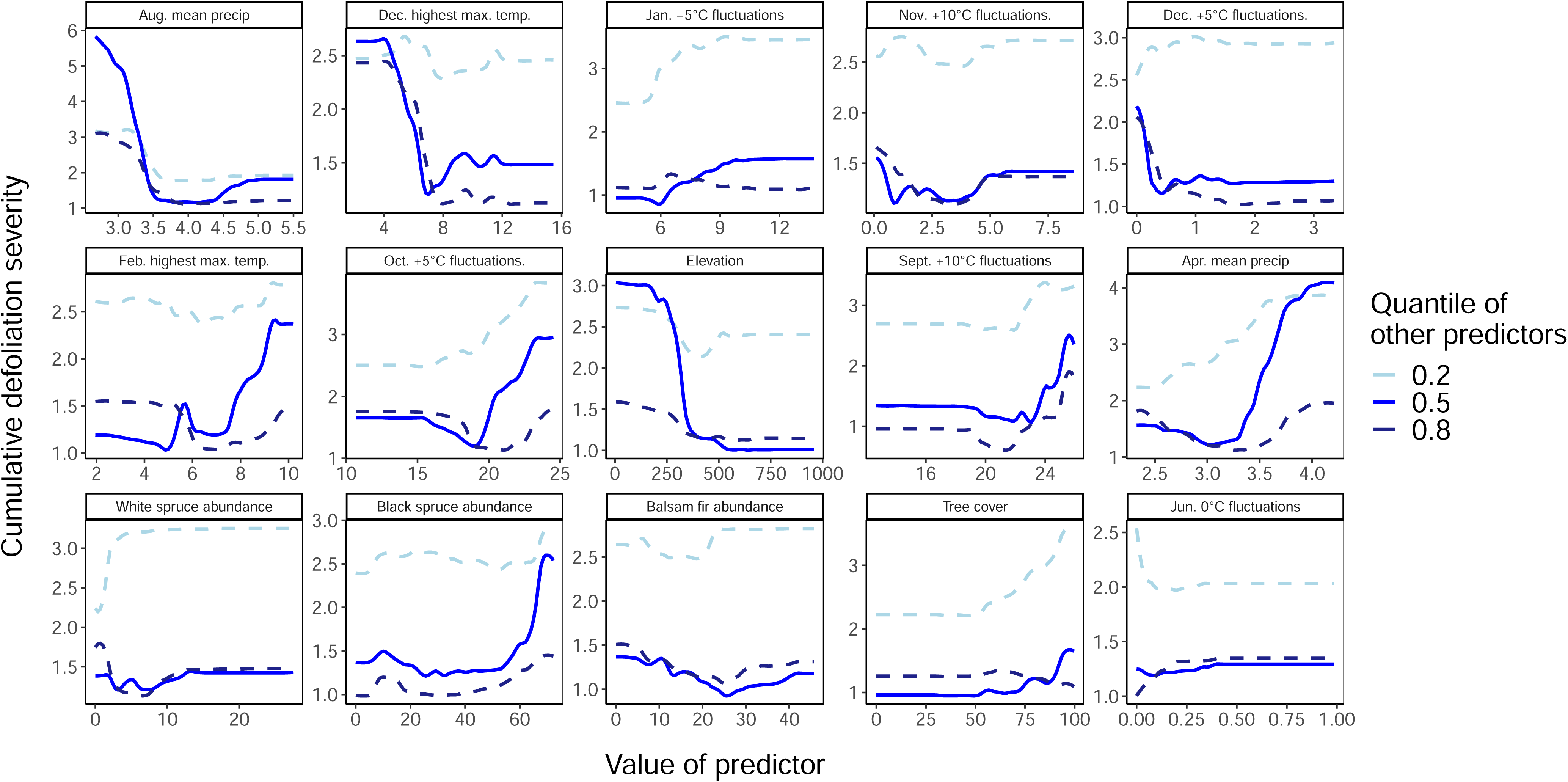
Relationship between random forest regression model predictors and predicted cumulative defoliation severity across fixed quantile values of other model predictors. X-axes indicates value of labelled predictor across outbreak range: temperatures = °C, fluctuations = frequency per month, tree abundances = % cover, elevation = m, and precipitation = kg m^-2^ s^-1^ (mean rate of precipitation scaled up by a factor of 100,000 to ensure compatibility with predictor selection).

### iii) Climate change predictions

In general, the regression models trained with and without temperature fluctuation were able to differentiate areas susceptible to defoliation when predicting using CMIP5 predicted climatologies for 2041-2070 (Figure 5A,B). The model trained with temperature fluctuation variables had an average predicted cumulative defoliation severity of 2.44 ± 0.68 (SD) and predicted the highest defoliation severity as 4.87. The model trained without temperature fluctuation variables predicted higher defoliation on average, with a mean predicted cumulative defoliation severity of 3.03 ± 0.64 (SD). This model also predicted a higher maximum cumulative defoliation severity, with a maximum predicted value of 6.70.

**Figure 5.**
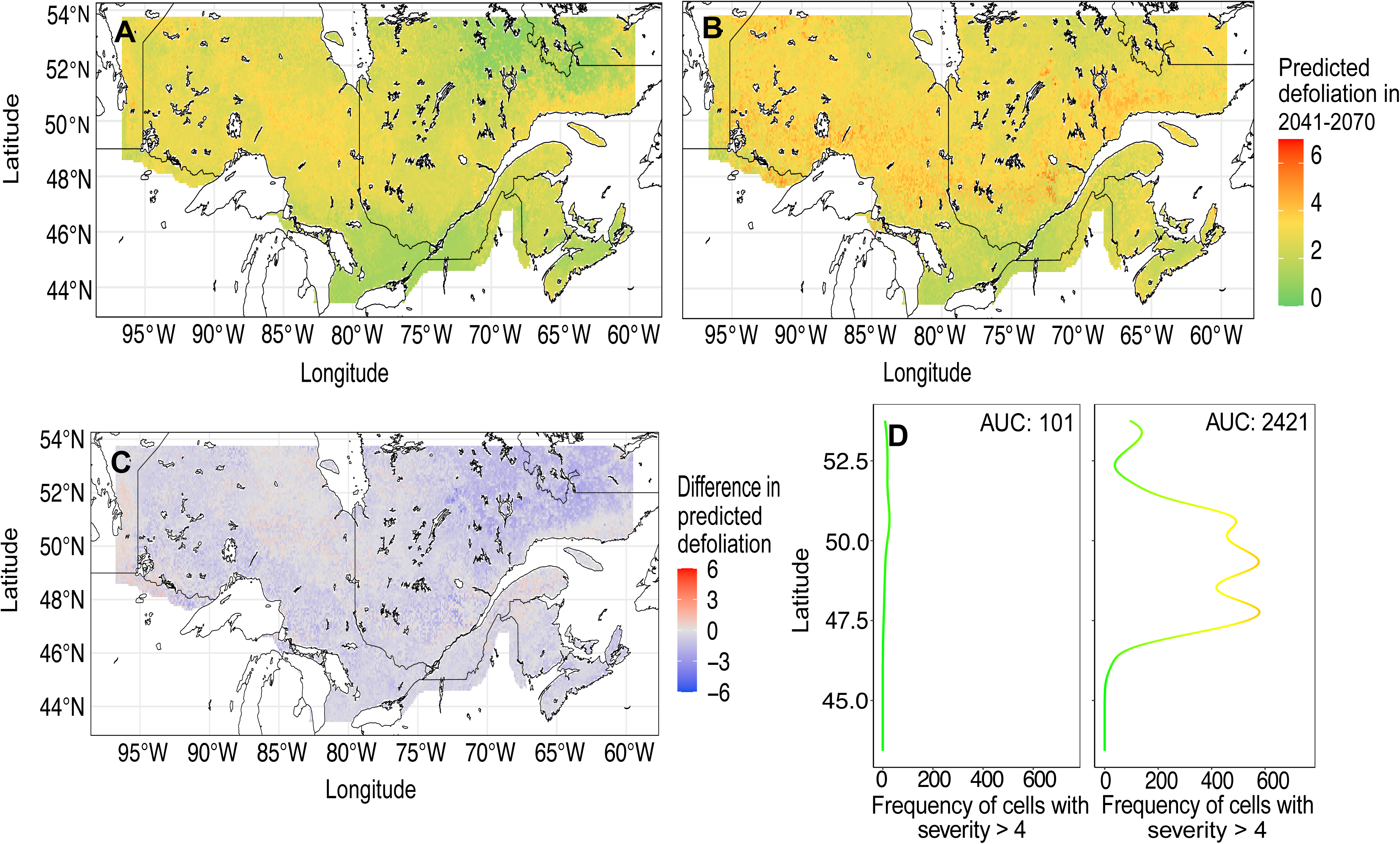
Predicted future defoliation across Quebec, Ontario, and Labrador by the random forest regression model using CMIP5 predicted climatologies for 2041-2070. Areas covered by lakes and rivers removed from prediction area. **A)** Distribution of predicted cumulative defoliation severity of model trained using temperature fluctuations. **B)** Distribution of predicted cumulative defoliation severity of model trained without temperature fluctuations. **C)** ΔPredicted defoliation severity between models trained with and without temperature fluctuations. **D)** The model trained using temperature fluctuations predicted severe (>4 cumulative severity score) defoliation less and with a more northerly distribution

The regression model trained with temperature fluctuations predicted severe defoliation at a lower frequency than the model trained without temperature fluctuations, but in more northerly regions (Figure 5). The frequency of cells with a predicted cumulative defoliation severity >4 was much higher in the model trained without temperature fluctuations (AUC_with fluctuations_ = 101; AUC_without fluctuations_ = 2,421). The average latitude of cells with a defoliation severity >4 was 51.1 ± 1.54 (SD) in the model trained with temperature fluctuations, and 49.4 ± 1.66 (SD) in the model without. However, given higher defoliation severity is observed near to the highest latitudes predicted on by the model it is possible that the latitudinal extent of the study area is artificially truncating defoliation predictions.

## Discussion

Here, we show the importance of including physiologically-informed predictors in species distribution models as we find temperature fluctuations are an important predictor of spruce budworm defoliation from 2006-2016 in Quebec. Including temperature fluctuations improved the performance of the random forest regression model and impacted predictions of future defoliation under climate change. Temperature fluctuation predictors ranked highly amongst model predictors and had a similar mean importance to climate predictors and higher mean importance than forest predictors. When predicting future defoliation under climate change conditions, the inclusion of temperature fluctuation predictors drastically reduced predicted defoliation severity across the landscape. These results suggest that temperature fluctuations play a role in influencing the landscape-scale defoliation of spruce budworm and demonstrate the importance of incorporating physiological measures into species distribution models of ectotherms to improve performance.

The importance of temperature fluctuations to the defoliation regression model was expected given the physiological importance of temperature fluctuations to spruce budworm in the laboratory and to other ectotherms generally. Marshall and Sinclair (2015) demonstrated that the number and frequency of low temperature stress events experienced affected spruce budworm survival in the laboratory, which provides experimental evidence that supports our findings that temperature fluctuations may be important on a landscape scale. Their results also indicate that the frequency of temperature fluctuations may be more important than duration and severity of temperature exposure, which was not the case in our model given the importance of other climatic variables. However, climate may matter more at large spatial scales than in laboratory experiments due to climate extremes in the northern and southern spruce budworm distribution margins that define range limits (Marshall & Roe, 2021). Additionally, our model suggests that fluctuations at mild temperatures during the fall and winter may influence defoliation severity over the landscape. How the frequency of mild temperature exposures may affect spruce budworm and other ectotherms during the fall and winter has not been directly studied. However, short exposures to warm temperature events are known to affect the fitness of spruce budworm and other insects, particularly when they occur prior to or during overwintering in the fall and winter. Roe et al. (2024) found that diapause preparation in spruce budworm is highly sensitive to fall length and temperature, and that there is a delicate balance between survival and energetic costs during this period. Prolonged exposure to higher temperatures prior to overwintering also led to significant delayed post-overwintering mortality in the green-veined white (*Pieris napi*; Nielsen et al., 2022). Thus, while there is no direct correlation between spruce budworm fitness and mild temperature fluctuations in fall and winter, it is plausible that such a relationship may exist, and we suggest further study to identify mechanistic explanations.

In general, the importance of climate predictors to the random forest regression model was consistent with findings from previous spruce budworm species distribution models (Candau & Fleming, 2005; Gray, 2008; Zhang et al., 2023) and environmental factors known to affect spruce budworm physiology and defoliation. Winter (Dec. and Feb.) maximum temperatures and spring and summer (Apr. and Aug.) precipitation were the most important climate predictors to our model. The importance of winter maximum temperatures is consistent with the spruce budworm distribution model created by Candau and Fleming (2005), who identified this as the most important variable for predicting defoliation frequency during a 1967-1998 outbreak in Ontario. Given the effect of winter temperatures on insect cold stress and overwintering survival and fitness (Marshall & Roe, 2021; Sinclair, 2015), it is logical that winter temperatures may also drive spruce budworm fitness at the landscape scale. Precipitation was also important for predicting defoliation; in particular, August precipitation was the most important individual predictor of defoliation in our regression model. Studies on spruce budworm show that periods of drought can promote defoliation (Wellington et al., 1950), and that larval development is faster in summers with dry and sunny weather (Greenbank, 1956); however, spruce budworm have generally eclosed by August so climatic conditions in this month likely do not have a direct effect on larval defoliation (Marshall & Roe, 2021). Peak spruce budworm moth exodus typically occurs in August (Royama, 1984), so lower tropospheric weather conditions in this month influence dispersal (Boulanger et al., 2017) and may partially explain our results. Our results match the spruce budworm defoliation model created by Zhang et al. (2023), as they also found that spring and summer precipitation are strong predictors of defoliation during an outbreak in Newfoundland. Ultimately, our results align with previous findings on the climatic drivers of defoliation and underscore the influence of climate on spruce budworm outbreak dynamics.

Our model provides insights into the factors influencing the distribution and severity of spruce budworm defoliation; while these insights are ultimately correlative, they are supported by physiological mechanisms and provide a case study for incorporating physiology into species distribution models. Correlative models use climate and environmental factors to predict the response of an organism to changes in those factors, and have been used effectively to investigate the drivers of landscape-scale spruce budworm outbreaks (Candau & Fleming, 2005; Gray, 2008; Zhang et al., 2023). However, these models do not consider organism physiology or sub-processes, which can modify the response of the organism to different conditions and inform their responses to climate change (Maino et al., 2016). Thus, robust predictions of ectotherm distributions in changing conditions require an understanding of physiological mechanisms driving their response in addition to landscape-level studies of abundance and fitness using modelling techniques (Rodríguez et al., 2019). Our model demonstrates that incorporating spatial predictors which directly relate to species survival and fitness is possible and improves species distribution model performance, and adds to a growing body of work on “hybrid” species distribution modelling which combine mechanistic and correlative approaches (Tourinho & Vale, 2023). These model principles may also be extended upon in future models by incorporating explicitly mechanistic methods such as *NicheMapR* (Kearney & Porter, 2020; Lembrechts et al., 2019) into correlative species distribution models.

Shifts in the frequency of temperature fluctuations under climate change may influence the future distribution of spruce budworm and other ectotherms, but challenges still exist when modelling future daily temperatures and how biota will respond to climate change. While the accuracy of species distribution model predictions generally cannot be evaluated, predicted spruce budworm defoliation under climate change by the model trained with temperature fluctuations was dramatically different than the model trained without and suggests that models are highly sensitive to these predictors. Overall temperature variability is expected to increase under climate change (Dillon et al., 2016; G. Wang & Dillon, 2014); however, the frequency of variation at specific temperature thresholds is strongly driven by warming fall, winter, and spring temperatures and the non-uniformity of climate change effects across biomes and latitudes (Contosta et al., 2024; Marshall et al., 2020). This was reflected in our results, as changes in the frequency of temperature fluctuation predictors under climate change varied widely across the landscape and among our predictors, as we saw a general decrease in the frequency of mild fall temperatures, and a general increase in the frequency of mild winter temperatures (unpublished data). This also highlights uncertainty in species distribution model predictions, as while our methods could generate stochastic daily temperatures informed by both daily historical temperatures and future climate trends, this may not accurately forecast trends in temperature fluctuations (and, by extension, spruce budworm defoliation and species distributions) in the future. More reliable methods to predict future daily maximum and minimum temperatures are being developed (Araya-Osses et al., 2020; Dillon et al., 2016; Requena et al., 2021), but there are still few resources on a fine spatial and temporal scale useful for large-scale biogeographical forecasting. The sensitivity of our species distribution model to temperature fluctuations and the lack of fine-scale future daily weather forecasting reveals uncertainty when predicting the response of biota to climate shifts. To combat this, we suggest further refinement of our methods and the development of more fine-scale future climate modelling to clarify the response of spruce budworm and other ectotherms to daily temperature fluctuations under climate change.

While our model attempts to capture how environmental factors influence spruce budworm defoliation, biotic factors may still explain the higher model error in areas of severe defoliation. The dynamics of spruce budworm outbreaks are complex and are likely driven by a combination of top-down and bottom-up factors (Pureswaran et al., 2016). Thus, higher model error in areas with high defoliation severity may be explained by spruce budworm population dynamics and defoliator-host tree relationships. De Grandpré et al. (2022) find that spruce budworm herbivory significantly increases tree needle nitrogen content, which may lead to a positive feedback loop of improved host foliage quality promoting further spruce budworm feeding and development. Additionally, spruce budworm population dynamics may account for the error distribution of our model. Dispersal of gravid moths can also synchronize outbreaks across the landscape (Régnière & Nealis, 2019), and population spread, growth rates, and outbreak risk are known to be density-dependent in the spruce budworm (Régnière et al., 2019). Thus, while models investigating environmental factors such as defoliation are useful, additional drivers contributing to outbreak dynamics that are not as easily modelled such as biotic interactions could be considered in future work.

### Conclusion

In this study, we investigated the hypothesis that inclusion of physiologically-informed predictors in a species distribution model improves performance using models of the landscape-scale distribution of spruce budworm defoliation. Results from our model of defoliation suggest temperature fluctuations influence historic spruce budworm defoliation in Quebec and are as important to the model as climatic variables. Additionally, inclusion of temperature fluctuation variables drastically decreases predicted defoliation severity across Ontario and Quebec under climate change conditions. Further study is needed to clarify the role of biotic drivers of outbreaks on defoliation patterns, and to generate reliable and large-scale future daily temperatures to more accurately evaluate the response of insects to climate shifts in the future. Ultimately, elucidating the effects of temperature fluctuations and other climatic factors on spruce budworm defoliation reveals a link between physiological responses and landscape-scale fitness that should be considered when building species distribution models of ectotherms. Given the highly destructive nature of spruce budworm, this model will contribute to our understanding of the drivers of forest pest outbreaks to enable better prediction of regions susceptible to large-scale outbreaks. Additionally, the effects of temperature fluctuations on survival and fitness are ubiquitous across ectotherm taxa; thus, our model provides a novel and useful framework for the incorporation of daily temperature variation into species distribution modelling.

## Supporting information

Supplemental methods, results, figures, tables

## Data availability statement

All data used in the study are publicly available and can be found in table S1. Code used in analyses, full-sized figures, supplementary material, and table S1 containing links to data are available at https://osf.io/anxc2/?view_only=aed3f40d549440eca89a38c0e12ab111.

## Acknowledgements

We would like to thank Dr. Amanda Roe, Ian DeMerchant, and Adam Hogg for their assistance in accessing and providing spruce budworm defoliation data, as well as the *Ministère des Ressources Naturelles et des Forêts* and the *National Forest Inventory* for data collection and curation. We would like to additionally thank Dr. Amanda Roe for her invaluable knowledge on spruce budworm and project guidance, and Dr. Allan Carroll for his guidance on modelling forest insect pests. We thank Dr. Amy Angert and Dr. Bill Milsom for their comments which significantly improved the preliminary manuscript.

## Citations

Alvich, J. (n.d.). Earth System (ESM4) (World). Retrieved October 5, 2023, from https://www.gfdl.noaa.gov/earth-system-esm4/

Araya-Osses, D., Casanueva, A., Román-Figueroa, C., Uribe, J. M., & Paneque, M. (2020). Climate change projections of temperature and precipitation in Chile based on statistical downscaling. Climate Dynamics, 54(9), 4309–4330. 10.1007/s00382-020-05231-4

Boisvert-Marsh, L., Pedlar, J. H., de Blois, S., Le Squin, A., Lawrence, K., McKenney, D. W., Williams, C., & Aubin, I. (2022). Migration-based simulations for Canadian trees show limited tracking of suitable climate under climate change. Diversity and Distributions, 28(11), 2330–2348. 10.1111/ddi.13630

Bouchard, M., & Auger, I. (2013). Influence of environmental factors and spatio-temporal covariates during the initial development of a spruce budworm outbreak. Landscape Ecology 2013 29:1, 29(1), 111–126. 10.1007/S10980-013-9966-X

Boulanger, Y., Fabry, F., Kilambi, A., Pureswaran, D. S., Sturtevant, B. R., & Saint-Amant, R. (2017). The use of weather surveillance radar and high-resolution three dimensional weather data to monitor a spruce budworm mass exodus flight. Agricultural and Forest Meteorology, *234–235*, 127–135. 10.1016/j.agrformet.2016.12.018

Candau, J. N., & Fleming, R. A. (2005). Landscape-scale spatial distribution of spruce budworm defoliation in relation to bioclimatic conditions. Canadian Journal of Forest Research, 35(9), 2218–2232. 10.1139/x05-078

Carrington, L. B., Armijos, M. V., Lambrechts, L., Barker, C. M., & Scott, T. W. (2013). Effects of fluctuating daily temperatures at critical thermal extremes on *Aedes aegypti* life-history traits. PLOS ONE, 8(3). 10.1371/JOURNAL.PONE.0058824

Colinet, H., Renault, D., Hance, T., & Vernon, P. (2006). The impact of fluctuating thermal regimes on the survival of a cold-exposed parasitic wasp, *Aphidius colemani*. Physiological Entomology, 31(3), 234–240. 10.1111/J.1365-3032.2006.00511.X

Contosta, A. R., Arndt, K. A., Baulch, H. M., Casson, N. J., Harpold, A., Morelli, T. L., Sirén, A. P. K., & Templer, P. H. (2024). Threshold Changes in Winter Temperature and Precipitation Drive Threshold Responses Across Nine Global Climate Zones and Associated Biomes. *Annual Review of Ecology*, Evolution, and Systematics, 55, 271–300. 10.1146/annurev-ecolsys-110421-102101

De Grandpré, L., Marchand, M., Kneeshaw, D. D., Paré, D., Boucher, D., Bourassa, S., Gervais, D., Simard, M., Griffin, J. M., & Pureswaran, D. S. (2022). Defoliation-induced changes in foliage quality may trigger broad-scale insect outbreaks. Communications Biology, 5(1), 1–10. 10.1038/s42003-022-03407-8

Dickerson, B. R., & and Vinyard, G. L. (1999). Effects of High Chronic Temperatures and Diel Temperature Cycles on the Survival and Growth of Lahontan Cutthroat Trout. Transactions of the American Fisheries Society, 128(3), 516–521. 10.1577/1548-8659(1999)128<0516:EOHCTA>2.0.CO;2

Dillon, M. E., Woods, H. A., Wang, G., Fey, S. B., Vasseur, D. A., Telemeco, R. S., Marshall, K., & Pincebourde, S. (2016). Life in the Frequency Domain: The Biological Impacts of Changes in Climate Variability at Multiple Time Scales. Integrative and Comparative Biology, 56(1), 14–30. 10.1093/ICB/ICW024

Dong, Y., Dong, S., Tian, X., Wang, F., & Zhang, M. (2006). Effects of diel temperature fluctuations on growth, oxygen consumption and proximate body composition in the sea cucumber *Apostichopus japonicus* Selenka. Aquaculture, 255(1), 514–521. 10.1016/j.aquaculture.2005.12.013

Dormann, C. F., Elith, J., Bacher, S., Buchmann, C., Carl, G., Carré, G., Marquéz, J. R. G., Gruber, B., Lafourcade, B., Leitão, P. J., Münkemüller, T., McClean, C., Osborne, P. E., Reineking, B., Schröder, B., Skidmore, A. K., Zurell, D., & Lautenbach, S. (2013). Collinearity: A review of methods to deal with it and a simulation study evaluating their performance. Ecography, 36(1), 27–46. 10.1111/j.1600-0587.2012.07348.x

Enriquez-Urzelai, U., Kearney, M. R., Nicieza, A. G., & Tingley, R. (2019). Integrating mechanistic and correlative niche models to unravel range-limiting processes in a temperate amphibian. Global Change Biology, 25(8), 2633–2647. 10.1111/gcb.14673

Findsen, A., Pedersen, T. H., Petersen, A. G., Nielsen, O. B., & Overgaard, J. (2014). Why do insects enter and recover from chill coma? Low temperature and high extracellular potassium compromise muscle function in *Locusta migratoria*. Journal of Experimental Biology, 217(8), 1297–1306. 10.1242/jeb.098442

González-Tokman, D., Córdoba-Aguilar, A., Dáttilo, W., Lira-Noriega, A., Sánchez-Guillén, R. A., & Villalobos, F. (2020). Insect responses to heat: Physiological mechanisms, evolution and ecological implications in a warming world. Biological Reviews, 95(3), 802–821. 10.1111/BRV.12588

Gray, D. R. (2008). The relationship between climate and outbreak characteristics of the spruce budworm in eastern Canada. Climatic Change, 87(3–4), 361–383. 10.1007/s10584-007-9317-5

Greenbank, D. O. (1956). The role of climate and dispersal in the initiation of outbreaks of the spruce budworm in New Brunswick: I. the role of climate. Canadian Journal of Zoology, 34(5), 453–476. 10.1139/z56-048

Hijmans, R. J., Bivand, R., Forner, K., Ooms, J., & Pebesma, E. (2021). *terra: Spatial Data Analysis* [Computer software]. Comprehensive R Archive Network (CRAN). https://CRAN.R-project.org/package=terra

Huang, L.-H., Chen, B., & Kang, L. (2007). Impact of mild temperature hardening on thermotolerance, fecundity, and Hsp gene expression in *Liriomyza huidobrensis*. Journal of Insect Physiology, 53(12), 1199–1205. 10.1016/j.jinsphys.2007.06.011

Johns, R. C., Bowden, J. J., Carleton, D. R., Cooke, B. J., Edwards, S., Emilson, E. J. S., James, P. M. A., Kneeshaw, D., MacLean, D. A., Martel, V., Moise, E. R. D., Mott, G. D., Norfolk, C. J., Owens, E., Pureswaran, D. S., Quiring, D. T., Régnière, J., Richard, B., & Stastny, M. (2019). A Conceptual Framework for the Spruce Budworm Early Intervention Strategy: Can Outbreaks be Stopped? Forests, 10(10), 910. 10.3390/F10100910

Karger, D. N., Conrad, O., Böhner, J., Kawohl, T., Kreft, H., Soria-Auza, R. W., Zimmermann, N. E., Linder, H. P., & Kessler, M. (2017). Climatologies at high resolution for the earth’s land surface areas. Scientific Data, 4(1), Article 1. 10.1038/sdata.2017.122

Karger, D. N., Wilson, A. M., Mahony, C., Zimmermann, N. E., & Jetz, W. (2021). Global daily 1 km land surface precipitation based on cloud cover-informed downscaling. Scientific Data, 8(1), 307. 10.1038/s41597-021-01084-6

Kearney, M., & Porter, W. (2009). Mechanistic niche modelling: Combining physiological and spatial data to predict species’ ranges. Ecology Letters, 12(4), 334–350. 10.1111/j.1461-0248.2008.01277.x

Kern, P., Cramp, R. L., & Franklin, C. E. (2015). Physiological responses of ectotherms to daily temperature variation. Journal of Experimental Biology, 218(19), 3068–3076. 10.1242/jeb.123166

Li, M., MacLean, D. A., Hennigar, C. R., & Ogilvie, J. (2020). Previous year outbreak conditions and spring climate predict spruce budworm population changes in the following year. Forest Ecology and Management, 458, 117737. 10.1016/J.FORECO.2019.117737

Luedeling, E., Caspersen, L., & Fernandez, E. (2023). *chillR: Statistical Methods for Phenology Analysis in Temperate Fruit Trees* (Version 0.75) [Computer software]. https://cran.r-project.org/web/packages/chillR/index.html

MacLean, D. A., Amirault, P., Amos-Binks, L., Carleton, D., Hennigar, C., Johns, R., & Régnière, J. (2019). Positive Results of an Early Intervention Strategy to Suppress a Spruce Budworm Outbreak after Five Years of Trials. Forests, 10(5), Article 5. 10.3390/f10050448

Maino, J. L., Kong, J. D., Hoffmann, A. A., Barton, M. G., Kearney, M. R., Koš, V., & Sinclair, B. J. (2016). Mechanistic models for predicting insect responses to climate change. Global Change Biology. 10.1016/j.cois.2016.07.006

Marshall, K. E., Gotthard, K., & Williams, C. M. (2020). Evolutionary impacts of winter climate change on insects. Current Opinion in Insect Science, 41, 54–62. 10.1016/J.COIS.2020.06.003

Marshall, K. E., & Roe, A. D. (2021). Surviving in a Frozen Forest: The Physiology of Eastern Spruce Budworm Overwintering. Physiology, 36(3), 174–182. 10.1152/PHYSIOL.00037.2020

Marshall, K. E., & Sinclair, B. J. (2010). Repeated stress exposure results in a survival-reproduction trade-off in *Drosophila melanogaster*. Proceedings of the Royal Society B: Biological Sciences, 277(1683), 963–969. 10.1098/rspb.2009.1807

Marshall, K. E., & Sinclair, B. J. (2012). The impacts of repeated cold exposure on insects. Journal of Experimental Biology, 215(10), 1607–1613. 10.1242/JEB.059956

Marshall, K. E., & Sinclair, B. J. (2015). The relative importance of number, duration and intensity of cold stress events in determining survival and energetics of an overwintering insect. Functional Ecology, 29(3), 357–366. 10.1111/1365-2435.12328

Mattson, W. J., Haack, R. A., Lawrence, R. K., & Slocum, S. S. (1991). Considering the nutritional ecology of the spruce budworm in its management. Forest Ecology and Management, 39, 183–210. 10.1016/0378-1127(91)90176-V

Ministère des Ressources naturelles et des Forêts. (2023). Aires infestées par la tordeuse des bourgeons de l’épinette au Québec en 2023 (p. 29).

Ministère des Ressources naturelles et des Forêts. (2024). *Aires infestées par la tordeuse des bourgeons de l’épinette au Québec en* 2024 (p. 34). Gouvernement du Québec.

National Forest Inventory. (2021). https://nfi.nfis.org/en

Natural Resources Canada. (2021). Spruce budworm. https://www.nrcan.gc.ca/our-natural-resources/forests/wildland-fires-insects-disturbances/top-forest-insects-and-diseases-canada/spruce-budworm/13383

Nielsen, M. E., Lehmann, P., & Gotthard, K. (2022). Longer and warmer prewinter periods reduce post-winter fitness in a diapausing insect. Functional Ecology, 36, 1151–1162. 10.1111/1365-2435.14037

Popp, A., Calvin, K., Fujimori, S., Havlik, P., Humpenöder, F., Stehfest, E., Bodirsky, B. L., Dietrich, J. P., Doelmann, J. C., Gusti, M., Hasegawa, T., Kyle, P., Obersteiner, M., Tabeau, A., Takahashi, K., Valin, H., Waldhoff, S., Weindl, I., Wise, M., … Vuuren, D. P. van. (2017). Land-use futures in the shared socio-economic pathways. Global Environmental Change, 42, 331–345. 10.1016/j.gloenvcha.2016.10.002

Pureswaran, D. S., Johns, R., Heard, S. B., & Quiring, D. (2016). Population Ecology Paradigms in Eastern Spruce Budworm (Lepidoptera: Tortricidae) Population Ecology: A Century of Debate Environmental Entomology Advance Access Downloaded from. Environmental Entomology, 1–10. 10.1093/ee/nvw103

Putnam, H. M., & Edmunds, P. J. (2011). The physiological response of reef corals to diel fluctuations in seawater temperature. Journal of Experimental Marine Biology and Ecology, 396(2), 216–223. 10.1016/j.jembe.2010.10.026

R Core Team. (2021). *R: A language and environment for statistical computing* [Computer software]. R Foundation for Statistical Computing.

Raynal, R. S., Noble, D. W. A., Riley, J. L., Senior, A. M., Warner, D. A., While, G. M., & Schwanz, L. E. (2022). Impact of fluctuating developmental temperatures on phenotypic traits in reptiles: A meta-analysis. Journal of Experimental Biology, 225(Suppl_1), jeb243369. 10.1242/jeb.243369

Régnière, J. (2009). Predicting insect continental distributions from species physiology. Unasylva, 60, 37–42.

Régnière, J., Cooke, B. J., Béchard, A., Dupont, A., & Therrien, P. (2019). Dynamics and Management of Rising Outbreak Spruce Budworm Populations. Forests, 10(9), Article 9. 10.3390/f10090748

Régnière, J., & Nealis, V. G. (2019). Density Dependence of Egg Recruitment and Moth Dispersal in Spruce Budworms. Forests, 10(8), Article 8. 10.3390/f10080706

Régnière, J., Powell, J., Bentz, B., & Nealis, V. (2012). Effects of temperature on development, survival and reproduction of insects: Experimental design, data analysis and modeling. Journal of Insect Physiology, 58(5), 634–647. 10.1016/J.JINSPHYS.2012.01.010

Régnière, J., St-Amant, R., & Duval, P. (2012). Predicting insect distributions under climate change from physiological responses: Spruce budworm as an example. Biological Invasions, 14(8), 1571–1586. 10.1007/S10530-010-9918-1/FIGURES/10

Requena, A. I., Nguyen, T.-H., Burn, D. H., Coulibaly, P., & Nguyen, V.-T.-V. (2021). A temporal downscaling approach for sub-daily gridded extreme rainfall intensity estimation under climate change. Journal of Hydrology: Regional Studies, 35, 100811. 10.1016/j.ejrh.2021.100811

Rodríguez, L., García, J. J., Carreño, F., & Martínez, B. (2019). Integration of physiological knowledge into hybrid species distribution modelling to improve forecast of distributional shifts of tropical corals. Diversity and Distributions, 25(5), 715–728. 10.1111/DDI.12883

Roe, A. D., Wardlaw, A. A., Butterson, S., & Marshall, K. E. (2024). Diapause survival requires a temperature-sensitive preparatory period. Current Research in Insect Science, 5, 100073. 10.1016/j.cris.2024.100073

Royama, T. (1984). Population Dynamics of the Spruce Budworm *Choristoneura fumiferana*. Ecological Monographs, 54(4), 429–462. 10.2307/1942595

Shiogama, H., Fujimori, S., Hasegawa, T., Hayashi, M., Hirabayashi, Y., Ogura, T., Iizumi, T., Takahashi, K., & Takemura, T. (2023). Important distinctiveness of SSP3–7.0 for use in impact assessments. Nature Climate Change, 13(12), 1276–1278. 10.1038/s41558-023-01883-2

Sinclair, B. J. (2015). Linking energetics and overwintering in temperate insects. Biology Publications, 72. https://ir.lib.uwo.ca/biologypubhttps://ir.lib.uwo.ca/biologypub/72

Sobek-Swant, S., Crosthwaite, J. C., Lyons, D. B., & Sinclair, B. J. (2012). Could phenotypic plasticity limit an invasive species? Incomplete reversibility of mid-winter deacclimation in emerald ash borer. Biological Invasions, 14(1), 115–125. 10.1007/S10530-011-9988-8/FIGURES/3

Srivastava, V., Tai, A. R., Robert, J. A., & Carroll, A. L. (2024). A dynamic outbreak distribution model (DODM) for an irruptive folivore: The western spruce budworm. Ecological Modelling, 492, 110737. 10.1016/j.ecolmodel.2024.110737

Storey, K. B., & Storey, J. M. (1989). Freeze Tolerance and Freeze Avoidance in Ectotherms. In L. C. H. Wang (Ed.), Animal Adaptation to Cold (pp. 51–82). Springer. 10.1007/978-3-642-74078-7_2

The SSP Scenarios. (n.d.). [Page]. DKRZ. Retrieved July 25, 2023, from https://www.dkrz.de/en/communication/climate-simulations/cmip6-en/the-ssp-scenarios

Tourinho, L., & Vale, M. M. (2023). Choosing among correlative, mechanistic, and hybrid models of species’ niche and distribution. Integrative Zoology, 18(1), 93–109. 10.1111/1749-4877.12618

Wang, G., & Dillon, M. E. (2014). Recent geographic convergence in diurnal and annual temperature cycling flattens global thermal profiles. Nature Climate Change, 4(11), Article 11. 10.1038/nclimate2378

Wang, T., Hamann, A., Spittlehouse, D., & Carroll, C. (2016). Locally Downscaled and Spatially Customizable Climate Data for Historical and Future Periods for North America. PLOS ONE, 11(6), e0156720. 10.1371/journal.pone.0156720

Wellington, W. G., Fettes, J. J., Belyea, R. M., & Turner, K. B. (1950). Physical and biological indicators of the development of outbreaks of the spruce budworm, *Choristoneura fumiferana* (clem.) (Lepidoptera: Tortricidae). Canadian Journal of Research, 28d(6), 308–331. 10.1139/cjr50d-021

Wright, M. N., & Ziegler, A. (2017). ranger: A Fast Implementation of Random Forests for High Dimensional Data in C++ and R. JSS Journal of Statistical Software, 77. 10.18637/jss.v077.i01

Zhang, B., Leroux, S. J., Bowden, J. J., Hargan, K. E., Hurford, A., & Moise, E. R. D. (2023). Species distribution model identifies influence of climatic constraints on severe defoliation at the leading edge of a native insect outbreak. Forest Ecology and Management, 544, 121166. 10.1016/j.foreco.2023.121166

