## Supplemental methods, results, figures, tables for "Temperature fluctuations influence predictions of landscape-scale spruce budworm defoliation in a physiologically-informed species distribution model"

*Ontario defoliation:*

Spruce budworm defoliation in Ontario is recorded by the Ministry of Natural Resources and Forestry during surveys of major insect pest damage in Ontario forests. Observed spruce budworm defoliation patches from 1996-2022 (n=39,866) were stored as polygons in a publicly available file containing records of all insect forest damage in Ontario (Ontario Ministry of Natural Resources and Forestry, 2022). The polygons have the following attributes: a unique numerical identifier, insect species, year of defoliation, forest damage ranking, notes, location accuracy, area of polygon, date of updates to polygon, and date recorded. The raster of spruce budworm defoliation in Ontario covered an area from -92.76°W to -74.98°W, and 42.87°N to 50.02°N (but was not used in subsequent analysis). Grids had an initial resolution of approximately 412m × 412m; however, grids used in analysis were aggregated to a resolution of 300 arcsec (approximately 6.5km × 6.5km) to match forest and climate predictors

Defoliation scores in Ontario were “light” (0-25% canopy damage), “moderate” (25-75%), “moderate-severe” (25-100% canopy damage), and “severe” (75-100%). To standardize data with Quebec records of spruce budworm defoliation, we filtered the dataset to only include spruce budworm records and assigned the defoliation level “light” a value of 1, the level “moderate” a value of 2, “moderate-severe” a value of 2.5 (as % defoliation spanned levels 2 and 3 in Quebec), and a value of 3 to “severe”. Cumulative defoliation severity across the landscape was calculated using the same methods as Quebec (Figure S1).

However, we encountered difficulties standardizing and analyzing records of defoliation in Ontario and Quebec. As well, the Quebec outbreak spanned a wider geographic area and was more severe; thus, we only used only records of defoliation from Quebec in modelling analyses.

***Supplemental citations***

Ontario Ministry of Natural Resources and Forestry. (2022, February 14). *Forest insect damage event*. https://www.arcgis.com/home/item.html?id=308957e79fce445fa74997c4cf501437

***Supplemental figures***

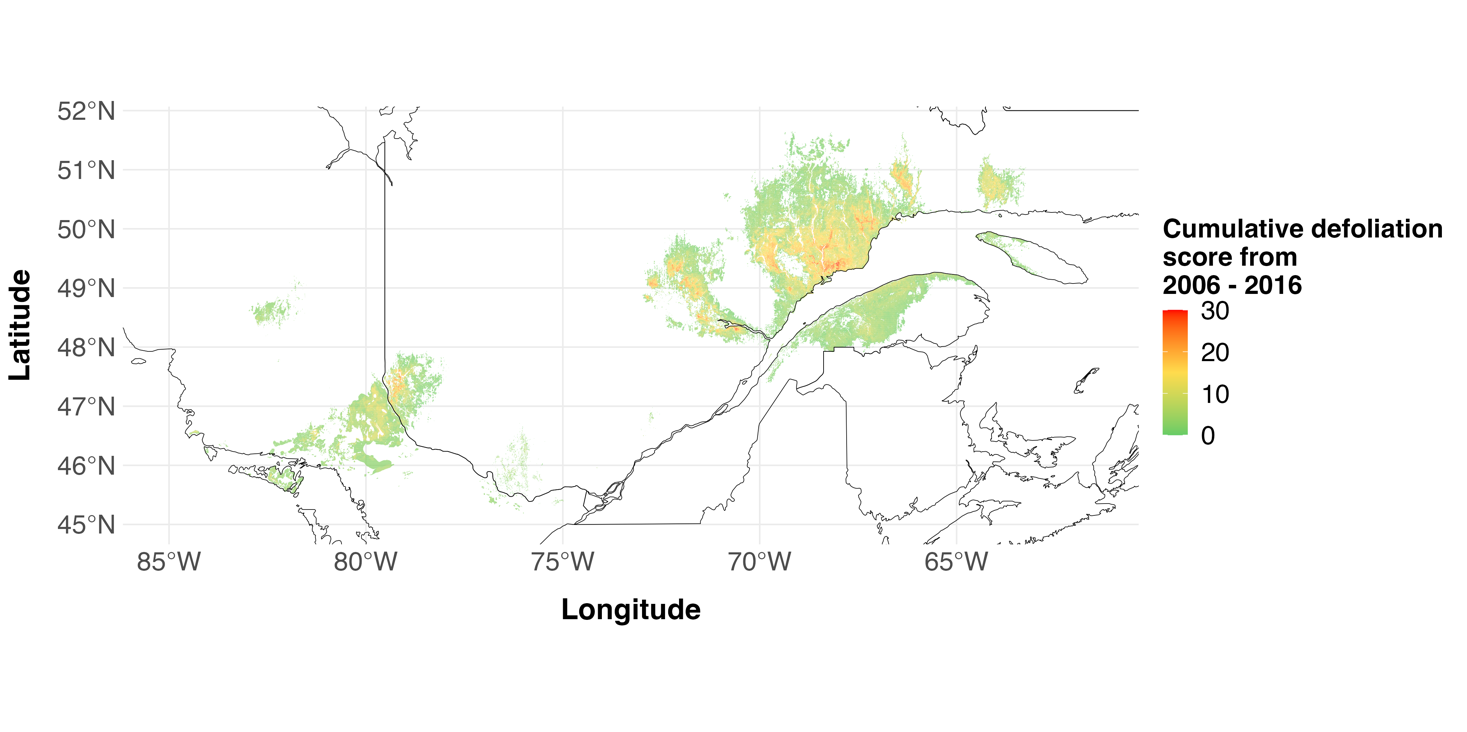

Figure S1. Spruce budworm outbreak distribution and severity in Quebec and Ontario from 2006 to 2016. Cumulative scores were generated by summing yearly defoliation polygon severity scores in each grid cell. Due to difficulties standardizing the data, only records from Quebec were used in modelling analyses. Provincial boundaries indicated by black lines. Resolution: 0.01 × 0.01 degrees latitude/longitude (~412 × 412m).

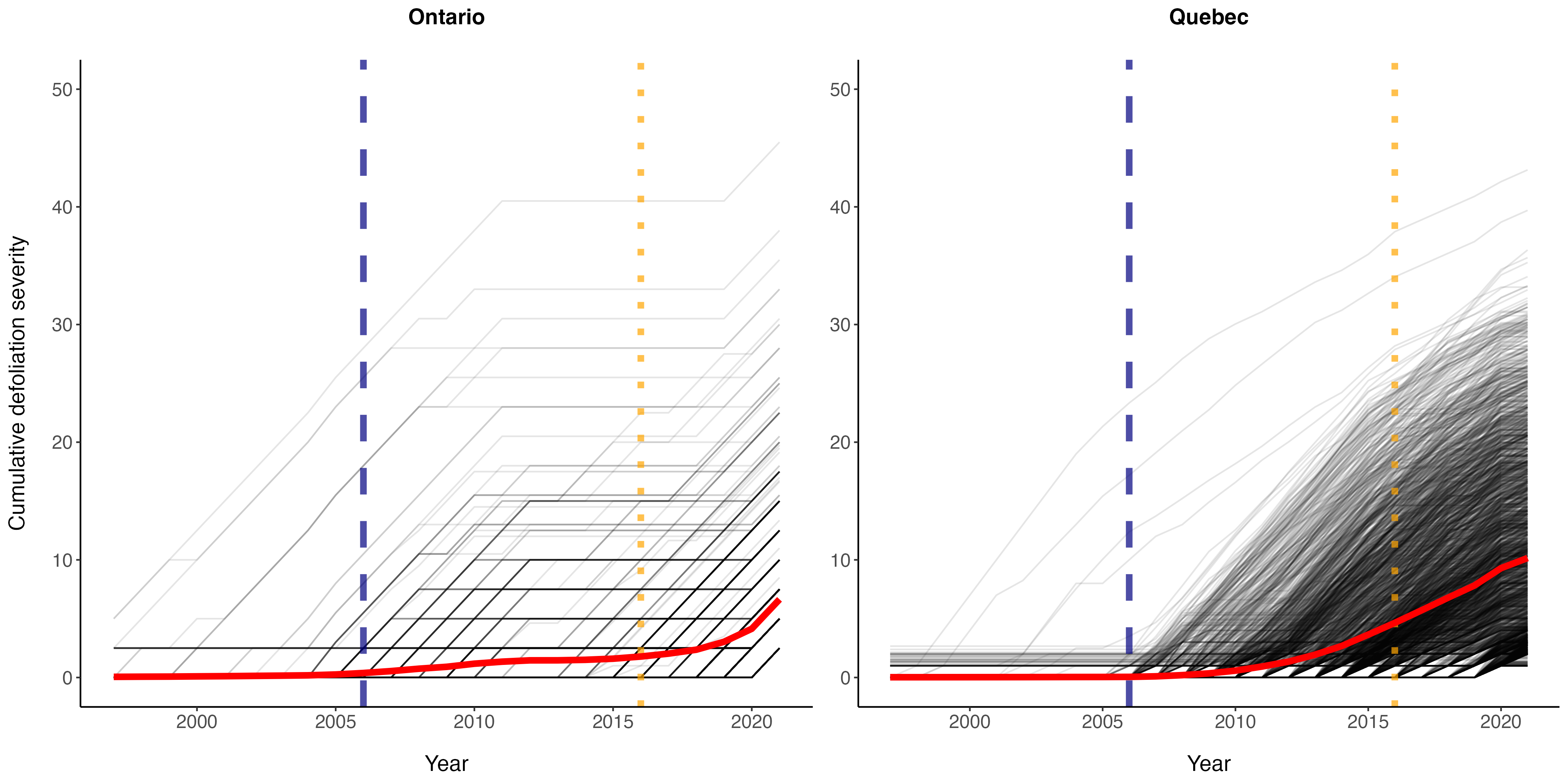

Figure S2. Cumulative defoliation severity of spruce budworm defoliation from 1997 - 2021 shows most recent outbreak in Ontario and Quebec beginning in 2006. Blue dashed line indicates year = 2006, orange dashed line indicates year = 2016. Black lines show cumulative defoliation severity observed in each grid cell across outbreak region. Red line indicates mean cumulative defoliation value across grid cells. Only points with non-zero defoliation in any year from 1992-2021 shown.

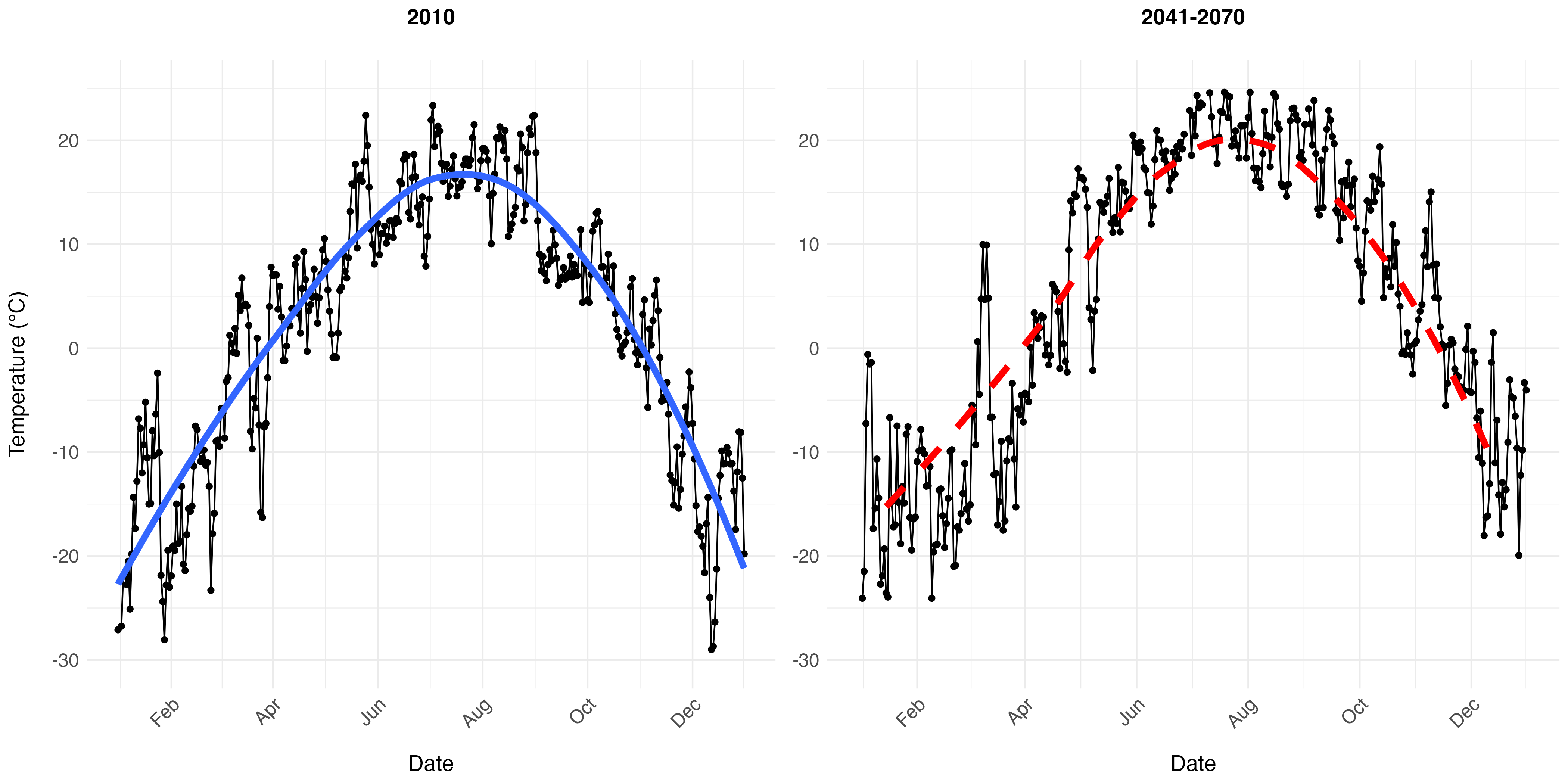

Figure S3. CHELSA monthly climate projections and the *chillR* package were used to generate daily stochastic temperatures that reflect trends in simulated climate change conditions. 2010 daily temperatures generated by CHELSA for a single grid cell (coordinates: -88.6°W, 50.9°N) are shown in comparison to daily temperatures generated for the same grid cell using monthly CHELSA climate projections and the *chillR* package. Black points and lines indicate daily minimum and maximum temperatures, red and blue lines show annual trends in temperature. Monthly CHELSA climate projections represent a single simulated year between 2041-2070.

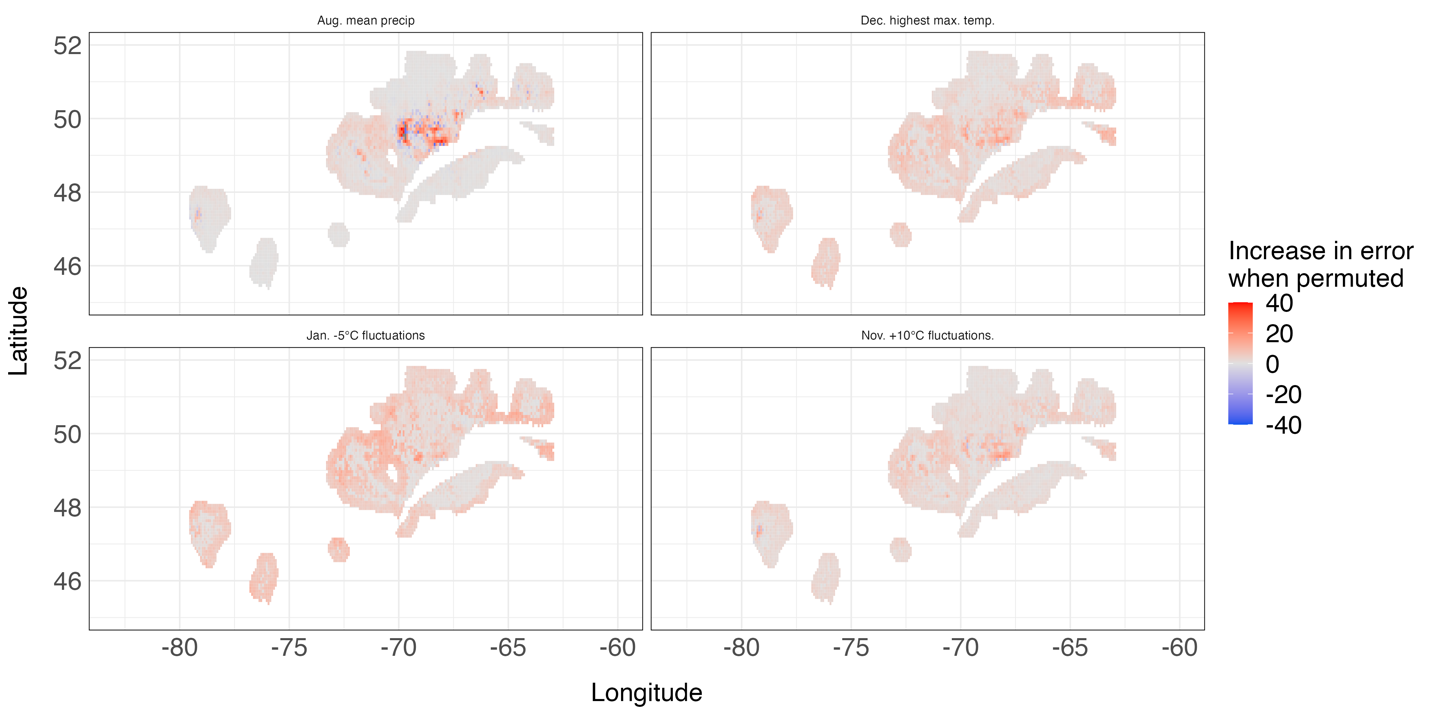

Figure S4. Local increase in error when top four most important variables to random forest regression model were permuted. Red colour indicates model performed worse when permuted, blue indicates model performed better.

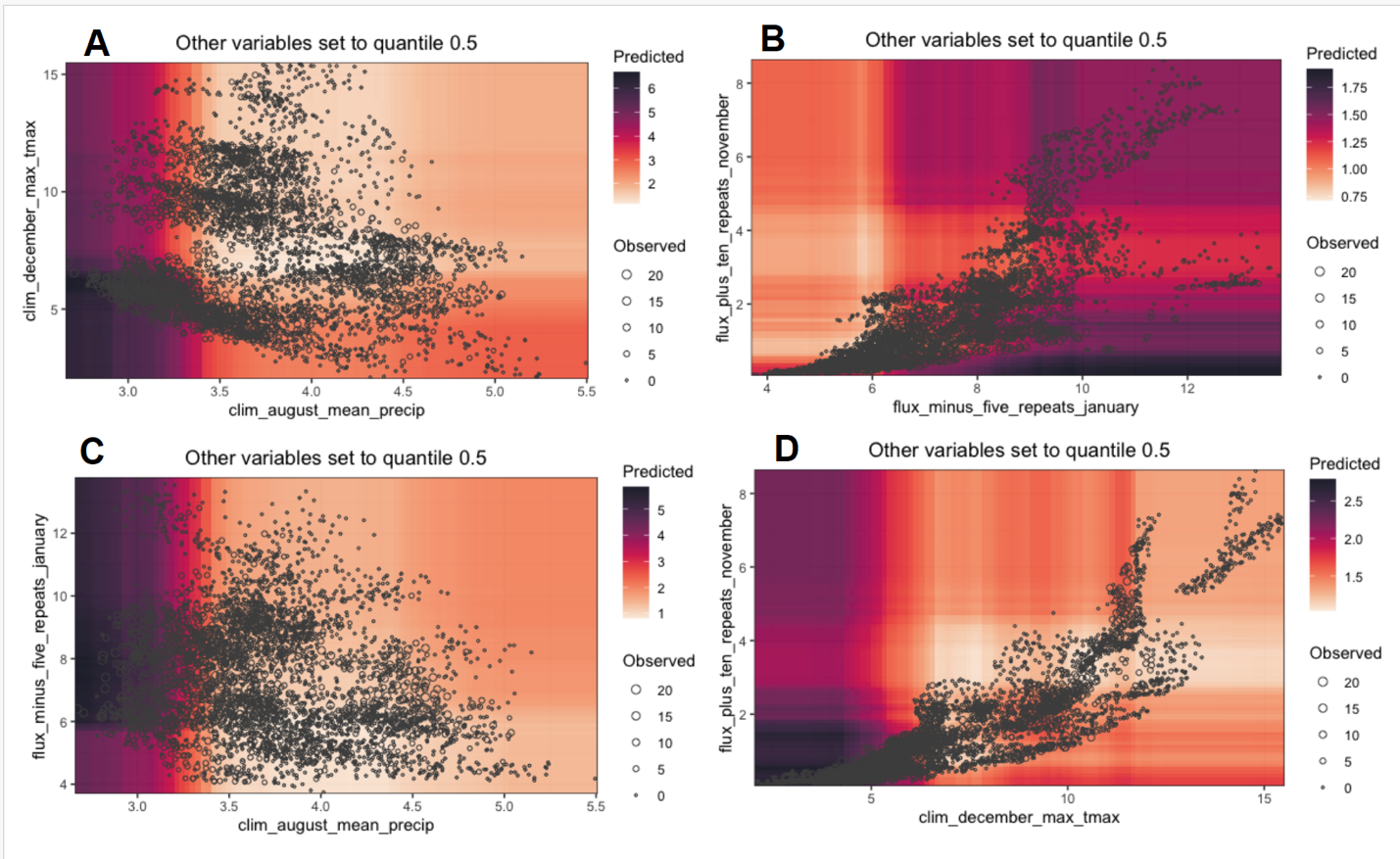

Figure S5. Response surface of top four most important predictors to random forest regression model. Other predictors set to quantile 0.5. Points (“Observed”) indicate observed defoliation severity, with size of point scaled with severity; colour gradient (“Predicted”) indicates predicted severity by regression model. **A)** August mean precipitation and December highest maximum temperature; **B)** January -5°C fluctuations and November +10°C fluctuations. **C)** August mean precipitation and January -5°C fluctuations. **D)** December highest maximum temperature and November +10°C fluctuations.

***Supplemental tables***

Table S1. Key resources table. Includes all resources used for research, their source, and a link to access data.

| **Resource** | **Source** | **URL** |
| --- | --- | --- |
| **Software** | | |
| R Version 4.3.1 | R Core Team | https://cran.r-project.org/bin/macosx/ |
| **Defoliation Data** | | |
| Quebec Defoliation, 2006-2016 | *Ministère des Ressources Naturelles et des Forêts* | https://diffusion.mffp.gouv.qc.ca/Diffusion/DonneeGratuite/Foret/PERTURBATIONS_NATURELLES/Tordeuse_bourgeons_epinette/ |
| Ontario Insect Damage Dataset | Ontario GeoHub | https://geohub.lio.gov.on.ca/documents/forest-insect-damage-event/about |
| **Predictor Data** | | |
| Forest Predictors | National Forest Inventory | https://ftp.maps.canada.ca/pub/nrcan_rncan/Forests_Foret/canada-forests-attributes_attributs-forests-canada/ |
| Climate/Temperature Fluctuation Predictors - Present | Climatologies at high resolution for the earth’s land surface areas  (CHELSA) | https://envicloud.wsl.ch/#/?prefix=chelsa%2Fchelsa_V2%2FGLOBAL%2F |
| Climate/Temperature Fluctuation Predictors - Future | Climatologies at high resolution for the earth’s land surface areas  (CHELSA) | https://envicloud.wsl.ch/#/?bucket=https%3A%2F%2Fos.zhdk.cloud.switch.ch%2Fenvicloud%2F&prefix=chelsa%2Fchelsa_V2%2FGLOBAL%2Fclimatologies%2F2041-2070%2FGFDL-ESM4%2F |
| Elevation | *Elevatr* package | https://registry.opendata.aws/terrain-tiles/;  https://opentopography.org/developers |

Table S2. List of predictors used for modelling, description of predictors, and number of predictors in each category.

| **Predictor** | **Description** | **Number of predictors** |
| --- | --- | --- |
| **Forest Predictors** | |  |
| Balsam fir cover (%) | % forest stand cover comprised of balsam fir | 1 |
| White spruce cover (%) | % forest stand cover comprised of white spruce | 1 |
| Black spruce cover (%) | % forest stand cover comprised of black spruce | 1 |
| Forest cover (%) | % of landscape covered with forest | 1 |
| **Climate Predictors** | |  |
| Mean annual temp (°C) | Mean of daily mean temperatures from Jan-Dec | 1 |
| Mean minimum annual temperature (°C) | Mean of daily minimum temperatures from Jan-Dec | 1 |
| Absolute minimum annual temperature (°C) | Absolute daily minimum temperature from Jan-Dec | 1 |
| Mean maximum annual temperature (°C) | Mean of daily maximum temperatures from Jan-Dec | 1 |
| Absolute maximum annual temperature (°C) | Absolute daily maximum temperature from Jan-Dec | 1 |
| Monthly mean temperature (°C) | Mean of daily mean temperatures in each month (Jan. – Dec.) | 12 |
| Monthly mean minimum temperature (°C) | Mean of daily minimum temperatures in each month (Jan. – Dec.) | 12 |
| Monthly absolute minimum temperature (°C) | Absolute daily minimum temperatures in each month (Jan. – Dec.) | 12 |
| Monthly mean maximum temperature (°C) | Mean of daily maximum temperatures in each month (Jan. – Dec.) | 12 |
| Monthly absolute maximum temperature (°C) | Absolute daily maximum temperatures in each month (Jan. – Dec.) | 12 |
| Precipitation (kg m^-2^ s^-1^) | Annual mean precipitation rate | 1 |
| Monthly precipitation (kg m^-2^ s^-1^) | Monthly mean precipitation rate (Jan. – Dec.) | 12 |
| Elevation (m) | Elevation | 1 |
| **Temperature Fluctuation Predictors** | |  |
| Temperature fluctuations at 25°C (frequency/month) | Number of times temperature crosses 25°C per month (May, Jun., Jul., Aug., Sept.) | 5 |
| Temperature fluctuations at 20°C (frequency/month) | Number of times temperature crosses 20°C per month (May, Jun., Jul., Aug., Sept., Oct.) | 6 |
| Temperature fluctuations at 15°C (frequency/month) | Number of times temperature crosses 15°C per month (Apr., May, Jun., Jul., Aug., Sept., Oct.) | 7 |
| Temperature fluctuations at 10°C (frequency/month) | Number of times temperature crosses 10°C per month (Apr., May, Jun., Jul., Aug., Sept., Oct., Nov.) | 8 |
| Temperature fluctuations at 5°C (frequency/month) | Number of times temperature crosses 5°C per month (Mar., Apr., May, Jun., Jul., Aug., Sept., Oct., Nov., Dec.) | 10 |
| Temperature fluctuations at 0°C (frequency/month) | Number of times temperature crosses 0°C per month (Jan., Feb., Mar., Apr., May, Jun., Sept., Oct., Nov., Dec.) | 10 |
| Temperature fluctuations at -5°C (frequency/month) | Number of times temperature crosses -5°C per month (Jan., Feb., Mar., Apr., May, Sept., Oct., Nov., Dec.) | 9 |
| Temperature fluctuations at -10°C (frequency/month) | Number of times temperature crosses -10°C per month (Jan., Feb., Mar., Apr., Oct., Nov., Dec.) | 7 |
| Temperature fluctuations at -15°C (frequency/month) | Number of times temperature crosses -15°C per month (Jan., Feb., Mar., Nov., Dec.) | 5 |
| Temperature fluctuations at -20°C (frequency/month) | Number of times temperature crosses -20°C per month (Jan., Feb., Dec.) | 3 |
| Temperature fluctuations at -25°C (frequency/month) | Number of times temperature crosses -25°C per month (Jan., Feb., Dec.) | 3 |
| **Degree Day Predictors** | | |
| Annual degree days (sum of difference between mean daily temperatures and 0°C) | Annual sum of differences between mean daily temperatures and 0°C | 1 |
| Monthly degree days (sum of difference between mean daily temperatures and 0°C) | Monthly sum of differences between mean daily temperatures and 0°C (Jan. – Dec.) | 12 |
| **Total:** | | 169 |

Table S3. Ranked predictor importance for defoliation models trained with and without temperature fluctuation predictors.

A) Predictor importance for model trained with temperature fluctuations.

| **Predictor category** | **Predictor** | **Increase in error when permuted** |
| --- | --- | --- |
| Climate | Aug. mean precip. | 2.227 |
| Climate | Dec. highest max. temp. | 2.119 |
| Fluctuations | Jan. −5°C fluctuations | 2.103 |
| Fluctuations | Nov. +10°C fluctuations | 2.093 |
| Fluctuations | Dec. +5°C fluctuations | 1.979 |
| Climate | Feb. highest max. temp | 1.874 |
| Fluctuations | Oct. +5°C fluctuations | 1.858 |
| Climate | Elevation | 1.813 |
| Fluctuations | Sept. +10°C fluctuations | 1.742 |
| Climate | Apr. mean precip | 1.725 |
| Forest | White spruce abundance | 1.637 |
| Forest | Black spruce abundance | 1.619 |
| Forest | Balsam fir abundance | 1.453 |
| Forest | Tree cover | 1.213 |
| Fluctuations | Jun. 0°C fluctuations | 1.177 |

B) Predictor importance for model trained without temperature fluctuations.

| **Predictor category** | **Predictor** | **Increase in error when permuted** |
| --- | --- | --- |
| Climate | Dec. highest max. temp. | 2.417 |
| Climate | Aug. mean precip. | 2.356 |
| Climate | Feb. mean max. temp. | 2.229 |
| Climate | Nov. highest max. temp. | 2.151 |
| Climate | Apr. highest max. temp. | 2.123 |
| Climate | Sept. lowest min. temp. | 2.001 |
| Climate | Apr. mean precip. | 1.922 |
| Forest | White spruce abundance | 1.83 |
| Forest | Black spruce abundance | 1.782 |
| Forest | Balsam fir abundance | 1.591 |
| Climate | Jun. mean precip. | 1.589 |
| Forest | Tree cover | 1.331 |
